# ClassiPhages 2.0: Sequence-based classification of phages using Artificial Neural Networks

**DOI:** 10.1101/558171

**Authors:** Cynthia Maria Chibani, Florentin Meinecke, Anton Farr, Sascha Dietrich, Heiko Liesegang

## Abstract

**Background/ Motivation:** In the era of affordable next generation sequencing technologies we are facing an exploding amount of new phage genome sequences. This requests high throughput phage classification tools that meet the standards of the International Committee on Taxonomy of Viruses (ICTV). However, an accurate prediction of phage taxonomic classification derived from phage sequences still poses a challenge due to the lack of performant taxonomic markers. Since machine learning methods have proved to be efficient for the classification of biological data we investigated how artificial neural networks perform on the task of phage taxonomy.

**Results:** In this work, 5,920 constructed and refined profile Hidden Markov Models (HMMs), derived from 8,721 phage sequences classified into 12 well known phage families, were used to scan phage proteome datasets. The resulting Phage Family-proteome to Phage-derived-HMMs scoring matrix was used to develop and train an Artificial Neural Network (ANN) to find patterns for phage classification into one of the phage families. Results show that using the 100 fold cross-validation test, the proposed method achieved an overall accuracy of 84.18 %. The ANN was tested on a set of unclassified phages and resulted in a taxonomic prediction. The ANN prediction was benchmarked against the prediction resulting of multi-HMM hits, and showed that the ANN performance is dependent on the quality of the input matrix.

**Conclusions:** We believe that, as long as some phage families on public databases are underrepresented, multi-HMM hits can be used as a classification method to populate those phage families, which in turn will improve the performance and accuracy of the ANN. We believe that the proposed method is an effective and promising method for phage classification. The good performance of the ANN and HMM based predictor indicates the efficiency of the method for phage classification, where we foresee its improvement with an increasing number of sequenced viral genomes.

## Introduction

Bacteriophages, bacterial viruses infecting bacteria, are of utmost importance due to the role they play in bacterial evolution (Roux et al. 2016). Virus classification is based on the idea of an evolutionary relationship between viruses and groups of viruses having more ability to exchange genetic material (Hans-W Ackermann 2011). Virus taxonomy is currently the responsibility of the International Committee on the Taxonomy of Viruses (ICTV). As of March 2017, there exist 4,404 approved Species, 735 Genera, 35 Subfamilies, 122 Families and 8 Orders (Lefkowitz et al. 2017).

The traditional method for the classification of phages is based on deciphering the type of nucleic acid and virion morphology using Transmission Electron Microscopy (TEM)(Rohwer & Edwards 2002). Experimental identification and classification of phages is based on physiological data and needs time to perform the experiments and expertise on the culture conditions of the corresponding host and phage system. However, within the explosive growth of phage sequences in the era of next generation sequencing technologies, there is an increasing amount of phage derived sequences that lack physiological data and knowledge on the host of the phages, especially in the case of metagenome data. This poses challenges to the successful implementation of a method which correctly classifies phages(Skewes-cox et al. 2014). Therefore, the development of a sequence based computational method, with the flexibility to integrate newly sequence derived phage descriptors, is necessary to allow rapid and accurate classification.

It is a known fact that phages do not have a ribosomal gene to place them on the tree of life (Rohwer & Edwards 2002). Phage classification based nucleotide pairwise comparison limits the process to similarities to phages found within reference databases (Bolduc et al. 2017). This poses a challenge to phage sequences identified from metagenomic datasets, where in one study by Paez-Espino et al (Paez-Espino et al. 2016), they identified over 125,000 contigs which revealed no sequence similarity to known viruses.

To that extent, taxonomic systems based on phage proteomes were suggested; however they come with their limitations (Meier-Kolthoff & Göker 2017). Clustering techniques optimized for viral classification were applied by Lima-Mendez et al. (Lima-Mendez et al. 2008)and Roux et al. (Roux et al. 2015), which showed the efficiency of the use of phage clustering as a basis of classification.

Profile HMMs proved to be a powerful method to model the sequence diversity of a set of orthologs, and thus are sensitive and more effective than pairwise alignment methods in detecting divergent viral sequences (Skewes-cox et al. 2014; Reyes et al. 2017). Additionally, Chibani et al. 2019 (accepted) showed that the use of a combination of phage derived profile HMM hits proved to be efficient to classify previously unclassified phage genomes into different phage families.

The emerging fields and use of machine learning and data mining in different biological fields are proving to be instrumental in answering challenging questions by looking into millions of biological data produced in the last decade. Because of their success with big data, ANNs and other machine learning models have gained a considerable amount of interest as a promising framework for biology. When combined with genomic information, novel machine learning and data mining techniques can advance the extraction of critical information and predict future observations from big data. Considerable progress has been made in the application of Support Vector Machines (SVM) (Manavalan, Tae H. Shin, et al. 2018; Tan et al. 2018) and Naïve Bayes (Feng et al. 2013) machine learning algorithms to identify phage virion proteins and in the application of ANN to classify tailed phages (currently deprecated) (Lopes et al. 2014). However, the use of machine learning for phage taxonomic classification has not been reported so far. Therefore, it is necessary to apply meaningful feature extraction and selection methods to investigate the classification method.

In order to address the limitations of current phage taxonomic classification software, we focused on the question of how profile HMMs (Chibani et al 2019 (accepted)) perform within a machine learning approach for the automated classification of phage genome sequences. We designed and developed an ANN, a well known supervised Machine Learning (ML) algorithm, which has been applied to several biological problems (Arango-Argoty et al. 2018; Seguritan et al. 2012). The ANN takes protein hits scores to phage derived profile HMMs per phage family as input, by applying a set of thresholds to select optimal features for a phage classification method. The performance of supervised prediction algorithms depends on the quality of the training data set. We therefore generated a training data set to train an ANN to classify new phage genomes and whether the public available phage genomes are sufficient. To our knowledge, this is the first ever reported use of ANN for the classification of phages into phage families with a trusted performance to accuracy ratio for the predictions.

## Materials and Methods

A five-step guideline has increasingly been endorsed (Manavalan, Tae Hwan Shin, et al. 2018) in a series of recent publications, to develop a sequence-based predictor for a biological system that can easily be used, which goes as follow:

(i) generating a solid benchmarking dataset to train and test the prediction model; (ii) formulate the biological sequence samples with an effective mathematical expression that can truly reflect their intrinsic correlation with the target to be predicted; (iii) develop a powerful algorithm to generate a prediction; (iv) implement cross-validation tests to objectively evaluate the performance of the predictor; and finally, (v) establish a user-friendly web-server for the predictor that is accessible to the public. Below, we describe the achieved steps.

## Data Collection

The raw phage dataset used in this research were retrieved from millardlab database (http://millardlab.org/bioinformatics/bacteriophage-genomes/).

As of 20 March 2018, the database contained in total 8,721 phage genomes (Table S1) belonging to 21 phage families summarized in **Table 1**.

**Table 1:**
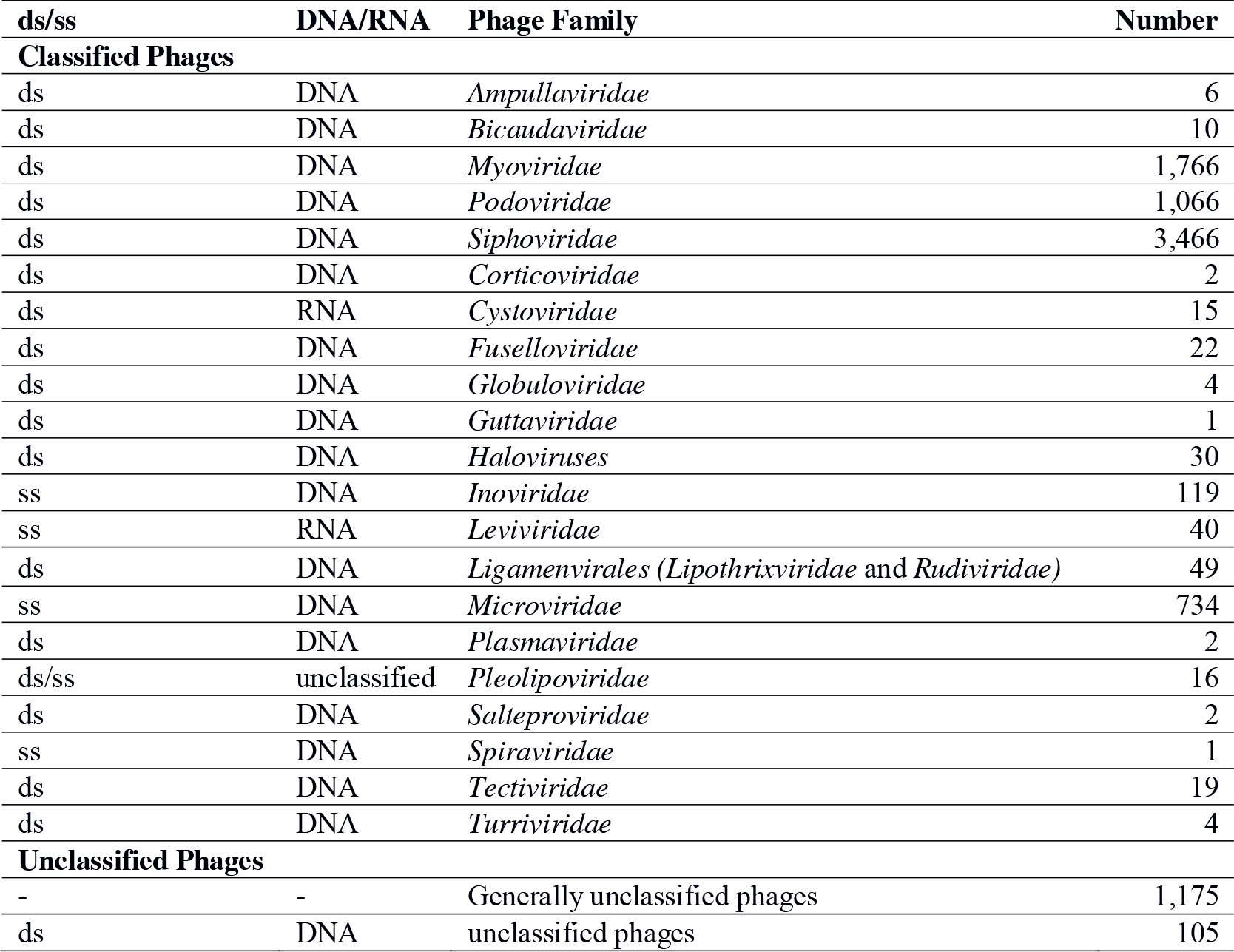
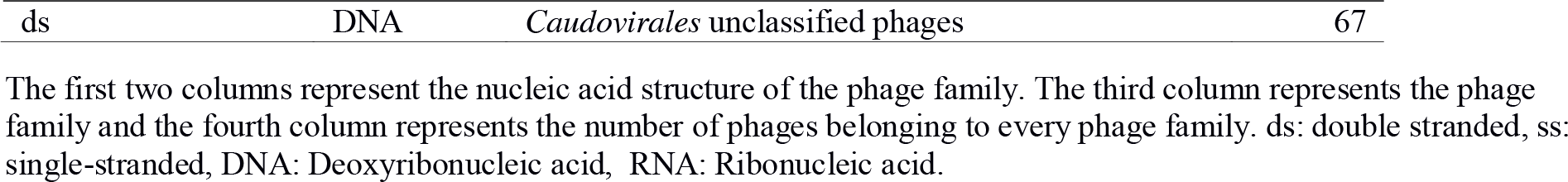
Summary table of the phage families and number of phages belonging to each phage family found in the millardlab database as of 20 March 2018

| ds/ss | DNA/RNA | Phage Family | Number |
| --- | --- | --- | --- |
| **Classified Phages** |  |  |  |
| ds | DNA | *Ampullaviridae* | 6 |
| ds | DNA | *Bicaudaviridae* | 10 |
| ds | DNA | *Myoviridae* | 1,766 |
| ds | DNA | *Podoviridae* | 1,066 |
| ds | DNA | *Siphoviridae* | 3,466 |
| ds | DNA | *Corticoviridae* | 2 |
| ds | RNA | *Cystoviridae* | 15 |
| ds | DNA | *Fuselloviridae* | 22 |
| ds | DNA | *Globuloviridae* | 4 |
| ds | DNA | *Guttaviridae* | 1 |
| ds | DNA | *Haloviruses* | 30 |
| ss | DNA | *Inoviridae* | 119 |
| ss | RNA | *Leviviridae* | 40 |
| ds | DNA | *Ligamenvirales (Lipothrixviridae and Rudiviridae)* | 49 |
| ss | DNA | *Microviridae* | 734 |
| ds | DNA | *Plasmaviridae* | 2 |
| ds/ss | unclassified | *Pleolipoviridae* | 16 |
| ds | DNA | *Salteproviridae* | 2 |
| ss | DNA | *Spiraviridae* | 1 |
| ds | DNA | *Tectiviridae* | 19 |
| ds | DNA | *Turriviridae* | 4 |
| **Unclassified Phages** |  |  |  |
| - | - | Generally unclassified phages | 1,175 |
| ds | DNA | unclassified phages | 105 |

The first two columns represent the nucleic acid structure of the phage family. The third column represents the phage family and the fourth column represents the number of phages belonging to every phage family. ds: double stranded, ss: single-stranded, DNA: Deoxyribonucleic acid, RNA: Ribonucleic acid.

## Data Construction

For the purpose of obtaining a reliable benchmark dataset, the following steps were considered. Phage families which had less than 15 phage genomes were excluded, in order to ensure diverse phages with diverse proteins for HMM generation. This step is crucial in order to differentiate between the highly biased number of *Siphoviridae* phages and least abundant ones. This resulted in 12 of the 21 phages families (*Cystoviridae*, *Fuselloviridae*, *Haloviruses*, *Inoviridae*, *Leviviridae*, *Ligamenvirales*, *Microviridae*, *Myoviridae*, *Pleolipoviridae*, *Podoviridae*, *Siphoviridae* and *Tectiviridae*) used for the benchmark dataset construction.

Non-redundant CDS, extracted from classified phage gbk files, were used as input for the Markov Clustering algorithm (MCL-edge). Clusters including more than 5 proteins were used to generate profile HMMs. Profile HMMs were subjected to refinement steps after rescanning the input extracted CDS. Refinement included 1) proteins not reaching the coverage threshold of 50% of the HMM consensus sequence were removed, and if were hit again, added to the model; 2) proteins removed due to redundancies were not added to the model; 3) proteins used to create the HMMs themselves if were hit were kept, if not hit thus were removed from the model; 4) not yet assigned proteins were added to the model. Rescanning the input and refinement steps were repeated until no change was observed. Resulting HMM scan bit-scores were normalized, and a set of input features were extracted, using the generated HMMs scanning the input data set, resulting in a cross-scan matrix of HMM-Phage-Family correlation to Protein-Phage correlation, we call Phage_input_matrix.

HMM profiles from the 12 phage families were generated as described by Chibani et al. 2019 (accepted) (see **Figure 1** for an overview of the methodology). In summary, protein coding sequences were extracted from the phage Gbk files, and sequences containing non-standard amino acid residues were excluded, as their meanings are ambiguous. To avoid biases and over-fitting, redundant proteins defined by CD-HIT (v.4.5.4)(Li & Godzik 2006) program by applying a 100% sequence identity cut-off, were removed during HMM generation steps. It should be noted that redundant proteins were removed only from the dataset used for HMM construction and not for the testing dataset. MCL-edge (v12-068) (Enright 2002) was used to generate protein clusters out of a BLASTp scan of all-against-all input protein sequences. For the clusters which had more than 5 proteins, multi-sequence alignment (MSA) files were generated. Profile HMMs were generated, per MSA file, using “hmmbuild” from HMMER (v3.1b1) (Finn et al. 2011) with default parameters. Removed proteins were stored for later refinement.

**Figure 1:**
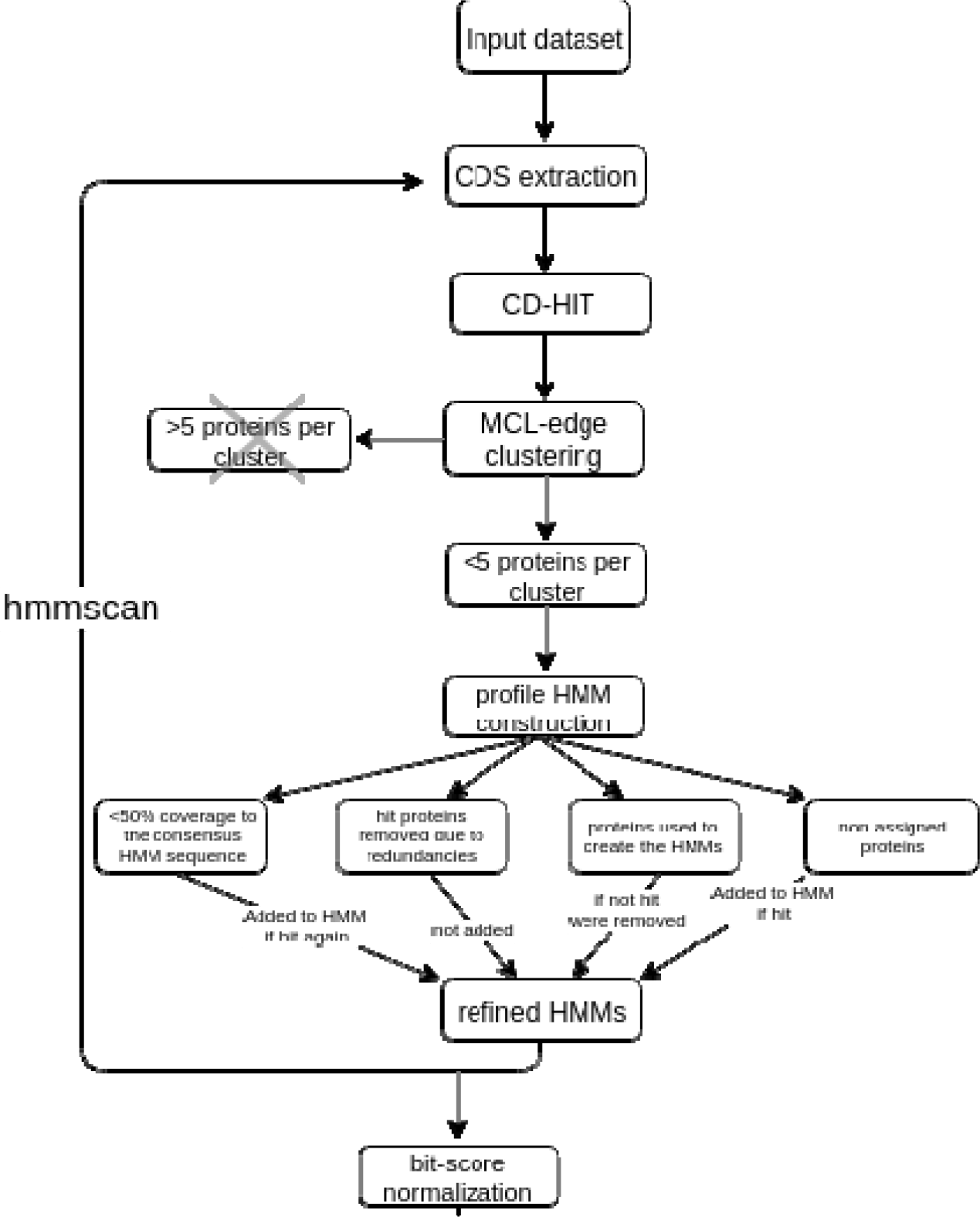
Overall framework of Phage_input_matrix construction.

The initially generated HMMs were then refined considering the following steps:

Firstly, the function “hmmemit” was used to create a consensus sequence from a generated profile HMM. This consensus sequence is closest in similarity to the majority of sequences used to create the respective HMM. Using “BLASTP” to align each protein of a cluster against the consensus sequence, proteins not reaching the coverage threshold of 50% were removed and stored for later refinement as well.

Secondly, the command “hmmpress” was used to create binary compressed data files (.h3m, .h3i, .h3f and .h3p) from a “profile HMM”. With “hmmscan” the binary files were used to look for orthologous protein hits in the scanned dataset. Created profile HMMs were used to scan the input fasta files where protein hits could be mapped to a) proteins removed due to redundancies b) proteins used to create the HMMs themselves c) not yet assigned proteins.

Lastly, proteins which are hit and have not yet been assigned were added to the profile HMM. Proteins that were used to create the HMM and were not hit, were removed from the profile HMM. Proteins that are hit but were previously removed due to redundancies were not added. Whenever multiple HMMs hit the same sets of proteins as well as their inputs, they were merged. Refined HMMs were used to rescan the input fasta and, if needed, refinement steps of merging were repeated until no changes occur. Resulting HMM scan bit-scores were lastly normalized (see Data normalization section) for further analysis.

## Feature extraction

The aim of this experiment was to train ANN Machine Learning (ML)-based model to accurately map input features generated from HMM scans, to predict the phage family a phage sequence belongs to, which is considered a multiclass classification problem. The key is to extract a set of informative features. We generated a set of input features for the ANN predictor, by scanning the proteomes of the 7,342 phages, of the remaining 12 phage families, using the generated 5,920 refined profile HMMs, which resulted in a cross-scan matrix of HMM-Phage-Family correlation to Protein-Phage correlation. The resulting bit-scores per HMM were extracted to generate input feature vectors for the training dataset with the phage family as the label.

For each individual phage of the phage family, one row is set up in the matrix, with the first two columns containing the bacteriophages name, which was later dropped, and phage family, which was used as the label. All other columns contain the bit-score value of the 5,920 HMM profiles scan of this phage protein sequences, or a default value of zero for no hit of that profile. We name our input matrix Phage_input_matrix.

## Data normalization

The bit-score values were normalized by dividing the resulting HMM scan bit-score by the number of amino acids of the consensus sequence of every HMM cluster. Hits of insufficient quality were filtered (e-score value <1e-10,(Amgarten et al. 2018; Arango-Argoty et al. 2018)). Additionally, if the bias of a hit was larger than the bit-score it produced, or if the bit-score was below zero in the first place, the corresponding HMM profile hit was omitted. If negative bit-score values were allowed, this would increase the value of empty hit cells in the final input matrix to a value greater than zero, creating values of HMM profile hits in the training dataset where there are none in the input.

After the creation of the matrix is completed and prior to the training of the ANN, its values are normalized to range from of [0,1], by employing “Minmax” formula described in (Manavalan et al. 2014):

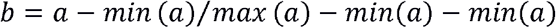

that can be used to reduce a k-dimensional array with any range to an array of the same shape covering a range from 0 to 1.

## Artificial Neural Network

We employed ANN as our algorithm, the objective of which is to learn to recognize patterns in a given dataset. Once it has been trained on samples of your data, it can make predictions by detecting similar patterns in future data(Schmidhuber 2015)). The “softMax” function (Manavalan, Tae H. Shin, et al. 2018), which is defined as *b = exp (ai)Σ exp(zj)*(Andrew Skabar, Dennis Wollersheim 2006), with a being a k-dimensional array. The resulting array, b, of the same shape as a, holds values ranging from 0 to 1 where all values in b add up to 1. Softmax was implemented as the activation function of the ANN’s output layers.

Based on the difference between the model’s predictions and the correct values, an error rate is calculated and the weights in each layer of the network are adjusted to reduce the error of the prediction. This procedure is performed from the output layer through the entire network to the input layer, hence the term back-propagation. The extent to which weights are adjusted is controlled by a learning rate. While linear and exponential decay functions did result in an increase of accuracy, the decay had to be gradual for the model to reach good prediction accuracy. This was achieved with high numbers of training epochs. We adapted the cosine decay, as discussed by (Loshchilov & Hutter 2016), proved to be the most efficient approach to decay the learning rate in our tested ANN architecture. In this study, we used the TensorFlow 1.10 package.

## Cross-Validation and Independent Testing

Usually, the benchmark dataset comprises a training dataset for training and a testing dataset for testing the model. Here, we performed 100-fold cross-validation on the training dataset and the trained model was tested on the independent dataset to confirm the generality of the developed method. For that, the benchmark dataset is split into 100 subsets, where 1/100^th^ of the initial data used for each of the testing subsets and the remainder used for training and cross-validation is performed using each of these 100 subsets as the testing dataset. The model trains for 100 individual sessions, once for each subset, as it must not have trained on any entry it later classifies in a testing set.

Here, all entries of the initial set are classified after the classification has ended, but the results can still vary due to the random distribution of entries in each training/testing subset. It should be noted that we performed 5 independent 100-fold cross-validations to confirm the robustness of the ML parameters.

## Performance Evaluation Criteria

To provide a simple method to measure the prediction quality, the following three metrics, sensitivity (Sn), specificity (Sp) and accuracy (Acc) were used and expressed as:

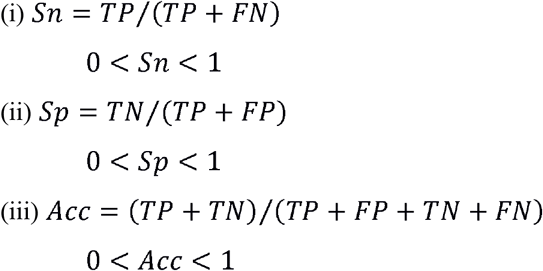

where TP is the number of phage correctly predicted to be of their corresponding phage families; TN is number of non-classified phages predicted to be not belonging to any phage family; FP in the number of is the number of non-classified phages predicted to belong to a phage family; and FN in the number of classified phages predicted not to belong to any phage family.

To further evaluate the performance of the ANN and determine suitable thresholds for the prediction values of the different families, we employed receiver operating characteristic (ROC) curves for the classification of each family. The ROC curve was plotted with the specificity as the x-axis and sensitivity as the y-axis by varying threshold. The area under the curve (AUC) was used for model evaluation, with higher AUC values corresponding to better performance of the classifier. The quality of the proposed method can be objectively evaluated by measuring the AUC.

## Results

### Data Construction

This method resulted in 5,920 refined profile HMMs, derived from 7,342 phages classified into 12 phage families (**Table 2**).

**Table 2:** Summary table of the number of refined HMMs resulting per phage family

| Phage Family | Refined HMMs |
| --- | --- |
| *Cystoviridae* | 2 |
| *Fuselloviridae* | 21 |
| *Haloviruses* | 48 |
| *Inoviridae* | 21 |
| *Leviviridae* | 4 |
| *Ligamenvirales* | 70 |
| *Microviridae* | 11 |
| *Myoviridae* | 2,851 |
| *Pleolipoviridae* | 3 |
| *Podoviridae* | 701 |
| *Siphoviridae* | 2,170 |
| *Tectiviridae* | 18 |

The first represents the phage family. The second column represents the number of refined HMMs generated per phage family.

The cross scan matrix resulting from the scan of HMMs derived from one phage family against the proteome of the 11 other phages families resulted in 60,560 protein hits by input HMM (Table S2).

### Neural Network Training and Classifications

The accuracy of the model during training was monitored using a scatter plot, which records the models performance on the testing set at every 10^th^ epoch of model training. Further collected metrics, the accuracy of the classification of the training and the testing data, as well as the learning rate at the given training epoch, were collected and plotted when training was complete (**Figure 2**). An overall prediction accuracy of 84.18 % was achieved by adopting ANN with a 100-fold cross-validation method on all phages in the dataset.

**Figure 2:**
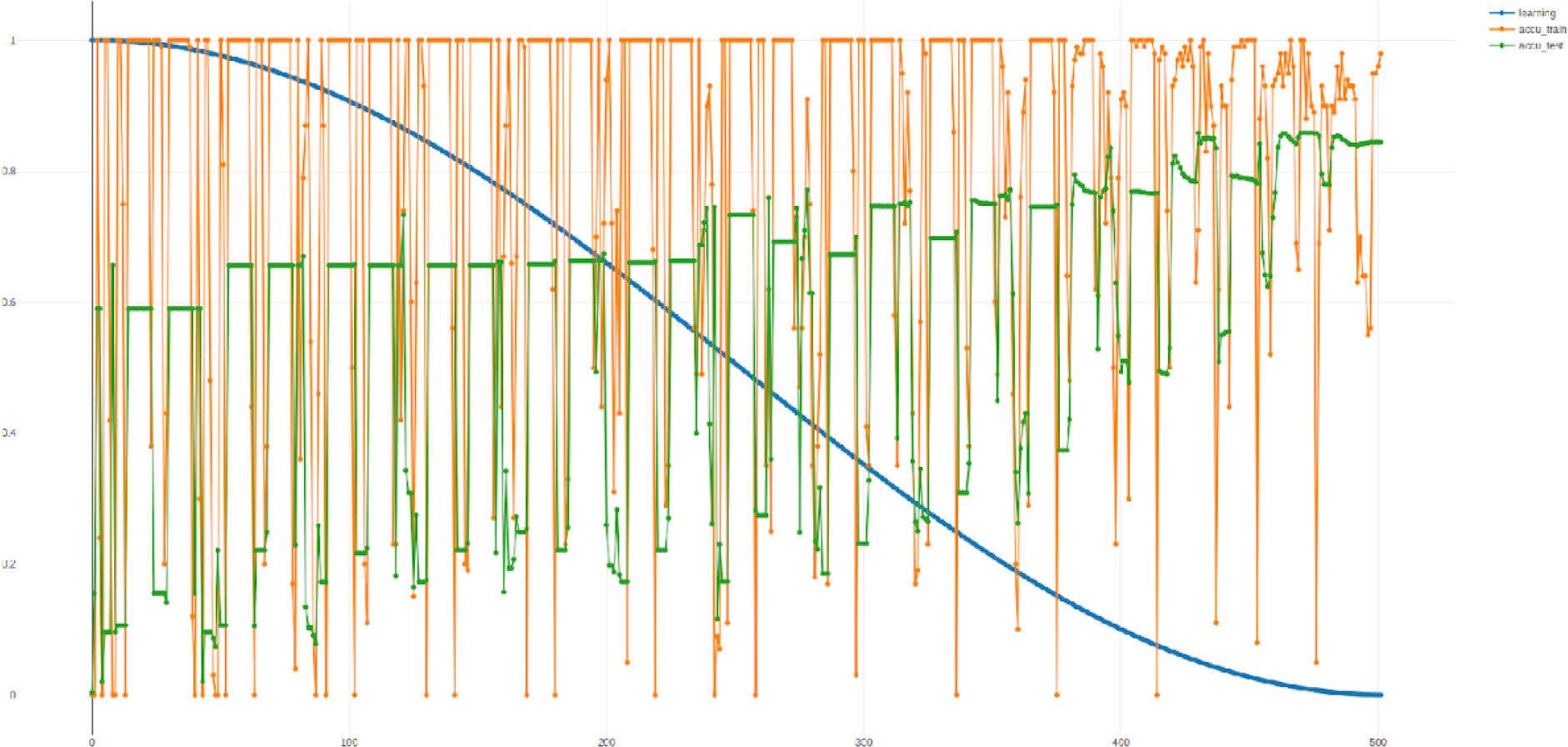
ANN performance on input matrix over training epochs. The plot displays the trends of the learning rate, training set accuracy and testing set accuracy over 500 epochs. The high learning rate in early epochs shows the high fluctuation of accuracies between epochs, as the adjustment of the model’s weights modifies it heavily. In the final epochs, the accuracy of the testing data classification reached 84.18%.

The scatter plot shows that the chosen batch size of 100 yielded the best result. We do not see information about possible issues with over- or under-fitting data. The model does not performs poorly on the testing set compared to the training set and thus did not result in over-fitting. Over-fitting results in a fluctuating training performance and low testing performance. Additionally, the model did not result in a poorer performance on both the training and the testing set. Under-fitting of the model to the training set results in a training performance curve that is constantly higher than the testing curve. The learning rate displays a decrease with an increasing number of epochs, to reach 0, when the accuracy of the testing reaches its high of 84.18%. We conclude there is no reason to assume issues with an over- or under-fitting model.

### Model performance and Metrics

The main output of the neural network is the label of the testing set and predictions of the model for each entry recorded at any training epoch. Using this information, the performance of the neural network can be accessed in detail for different stages of training. The labels of testing data are compared to the models assignments of the last recorded prediction by taking the maximum value of the models assignments.

As shown in **Table 3**, the TP, TN, FP, FN, Sp, Sn and Acc were calculated for the classification into the different phage families by using all 5,920 features.

**Table 3:** Predictive performance of the ANN per phage family

| Phage Family | TP | TN | FP | FN | Sensitivity | Specificity | Accuracy |
| --- | --- | --- | --- | --- | --- | --- | --- |
| *Cystoviridae* | 0 | 7,790 | 0 | 22 | 0 | 1 | 0.9971838 |
| *Fuselloviridae* | 0 | 7,782 | 0 | 15 | 0 | 1 | 0.9980762 |
| *Haloviruses* | 4 | 7,782 | 0 | 25 | 0.137931 | 1 | 0.9967994 |
| *Inoviridae* | 88 | 7,633 | 0 | 91 | 0.4916201 | 1 | 0.9883513 |
| *Leviviridae* | 25 | 7,776 | 0 | 11 | 0.6944444 | 1 | 0.9985919 |
| *Ligamenvirales* | 8 | 7,742 | 0 | 35 | 0.1860465 | 1 | 0.9955042 |
| *Microviridae* | 59 | 7,057 | 13 | 173 | 0.2543103 | 0.9981612 | 0.9745275 |
| *Myoviridae* | 577 | 6,548 | 40 | 647 | 0.4714052 | 0.9939284 | 0.9120584 |
| *Pleolipoviridae* | 0 | 7,796 | 0 | 16 | 0 | 1 | 0.9979519 |
| *Podoviridae* | 605 | 7,001 | 21 | 185 | 0.7658228 | 0.9970094 | 0.9736303 |
| *Siphoviridae* | 3,693 | 2,691 | 214 | 944 | 0.7964201 | 0.9325984 | 0.8517665 |
| *Tectiviridae* | 3 | 7,776 | 0 | 25 | 0.1071429 | 1 | 0.9967965 |

True or wrong phage classification prediction was assumed when the taxonomic prediction matched or did not match respectively the taxon that was given by the authors of the genome sequence. The number of correctly predicted phages (TP) of *Siphoviridae* (79.6%), *Podoviridae* (76.6%), *Leviviridae (*69.4 %), *Inoviridae* (49.1%), Myoviridae (45,5%), *Microviridae (*25.4%), *Haloviruses* (13.79%), *Ligamenvirales* (18.6%) and *Tectiviridae* (10.71%). Neither *Cystoviridae*, nor *Fuselloviridae*, or *Pleolipoviridae* were correctly predicted (TP = 0).

On the other hand, phage families where FP was predicted were *Microviridae*, *Myoviridae*, *Podoviridae* and *Siphoviridae*. All four phage families are known to infect bacterial hosts, however *Microviridae* are ss/DNA phages, whereas *Myo-*, *Podo*- and *Sipho-* are ds/DNA tailed phages belonging to the order of *Caudovirales*.

The clearest trend is the misclassification of entries to the *Siphoviridae* family. This occurs in families that are closely related to *Siphoviridae* (*Myoviridae*, *Podoviridae*), but also in structurally very distinct families such as *Fuselloviridae* and *Inoviridae*. This could indicate unexpected gene flux between unrelated phage species (Shapiro & Putonti 2018).

### ROC curves and thresholds

It is important to note that the confidence values in the final output of the model are not a percentage of likelihood for the corresponding entry. For example, a value of 0.7 as the highest value for an entry does not mean that the classification has a probability of 70% to be true. However, it makes it possible to set a threshold value to distinguish between more and less significant predictions. A higher threshold can improve the specificity of classification while a lower threshold results in highly sensitive classification. One threshold may have different effects on families, as the prediction scores are not calibrated between them. Thus, one score may be suited to distinguish true positives from false positives in one family but inappropriate to do this in another (Fawcett 2006). To determine suitable thresholds for the prediction values of different families, ROC curves for the classification of each family were created and plotted using the R package pROC (**Figure 3**).

**Figure 3:**
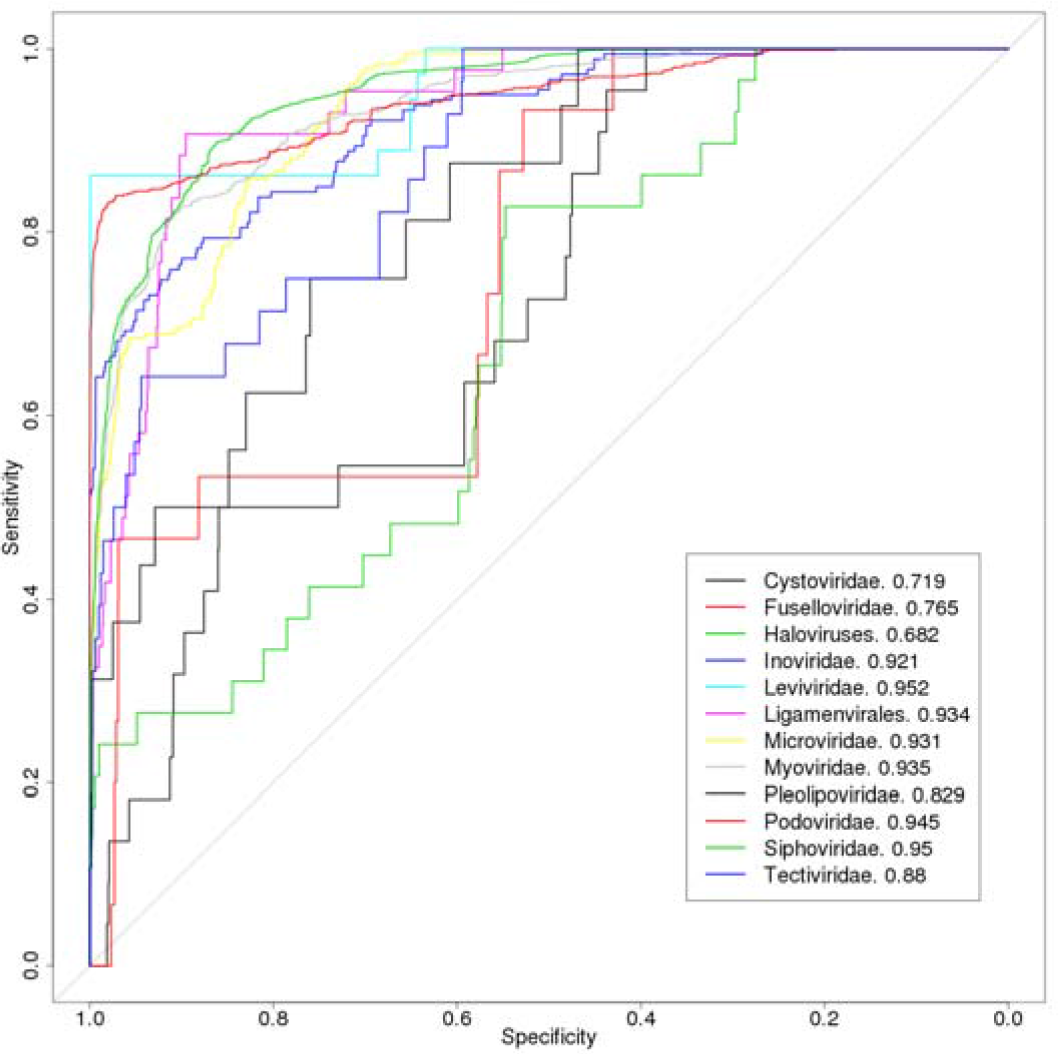
ROC curve resulting from the ANN classification.

ROC curves out of the input matrix dataset prediction. The performance of the neural network ranges from near perfect prediction (AUC of 0.97 for the *Leviviridae* family) to almost random (AUC of 0.682 for the *Pleolipoviridae* family). The varying trends of the individual curves reflect that classifications of different families benefit from thresholds that are unique to them

From the ROC curves, AUC (Area Under the Curve) values were calculated, which provided insight into the prediction performance without a specific threshold. As the area in a ROC plot is always 1, the area under the curve can range from 0 to 1, with 0.5 representing no predictive power and 1 perfect prediction. It can be interpreted as an average performance metric for the classifier. All calculated AUCs for were displayed in the legend of the ROC curves (AUC of 0.719 for *Cystoviridae*, 0.765 for *Fuselloviridae*, 0.682 for *Haloviruses*, 0.921 for *Inoviridae*, 0.952 for *Leviviridae*, 0.934 for *Ligamenvirales*, 0.931 for *Microviridae*, 0.935 for *Myoviridae*, 0.829 for *Pleolipoviridae*, 0.945 for *Podoviridae*, 0.95 for *Siphoviridae* and 0.88 for *Tectiviridae*).

### External dataset test

The proteomes of (∼1,347) unclassified phages (Generally unclassified phages, ds/DNA unclassified phages and ds/DNA/*Caudovirales* unclassified phages) were scanned using the set of 5,920 refined profile HMMs. A matrix using the resulting bit-scores per HMM was generated, where the bit-scores were normalized as was described previously. We used the generated ANN to test the ability of the ClassiPhage 2.0 model to predict the phage family classification of the unclassified phages. Out of 1,175 generally unclassified phages, predicted phage families were *Inoviridae*, *Microviridae*, *Myoviridae*, *Pleolipoviridae*, *Podoviridae*, *Siphoviridae* and *Tectiviridae*. Out of 105 ds/DNA unclassified phages, predicted phage families were *Microviridae*, *Myoviridae*, *Podoviridae*, *Siphoviridae* and *Tectiviridae*. Finally, out of 67 ds/DNA/*Caudovirales* unclassified phages, predicted phage families were *Halovirus*, *Microviridae*, *Myoviridae*, *Podoviridae* and *Siphoviridae* (Table S8). *Haloviruses* and *Microviridae* can’t be a classification for ds/DNA/*Caudovirales*, which shows that ClassiPhage 2.0 misclassifies phages where cross hits occur and enough family specific HMM hits.

We generate a heatmap of the prediction of the same set of unclassified vibriophages classified by Chibani et al 2019 (accepted) (**Figure 4**).

**Figure 4:**
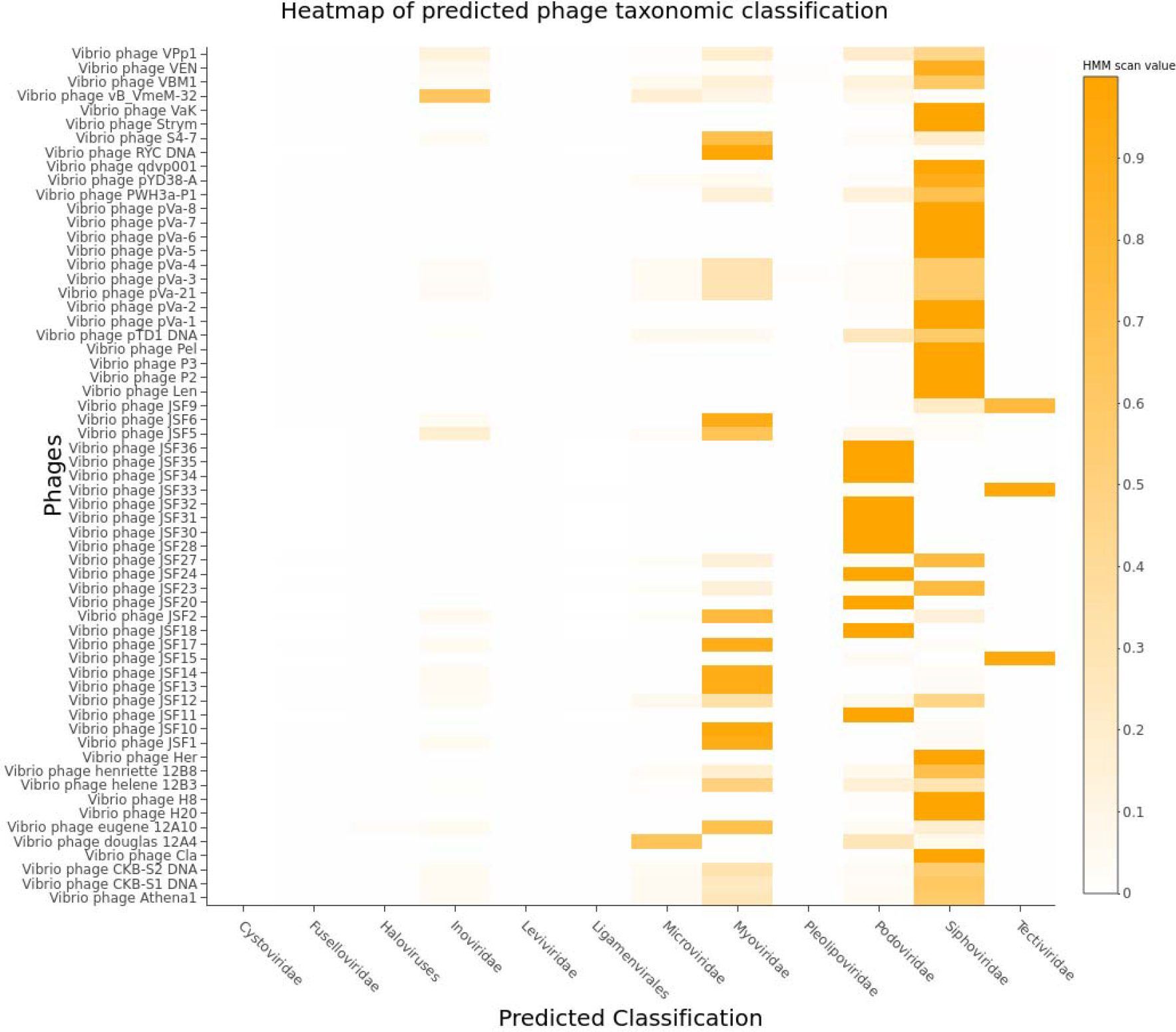
Heatmap of ClassiPhage 2.0 prediction of unclassified vibriophages. A heatmap based on a phage family prediction of a set of unclassified vibriophages by the ClassiPhage 2.0 model, displaying the phage labels (y-axis) and phage family prediction (x-axis).

22 classified phages were consistent with the classification resulting in Chibani et al. 2019 (accepted). 23 phages which had an unclear classification were classified as *Siphoviridae* by ClassiPhage 2.0. Lastly, out of 17 phages which were not consistent between the two methods, the clearest trend was the misclassification of entries to the *Siphoviridae* phage family (Table S9).

### Comparison to other methods

To the best of our knowledge, there exists no theoretical method for phage classification into phage families. Therefore, we cannot provide the comparison to analysis with published results to confirm that the model proposed here is superior to other methods. However, we generated a matrix out of the expected phage classification, as described in Chibani et al 2019 (accepted), to which we compare the prediction of ClassiPhage 2.0 of the unclassified dataset. We display phage predictions resulting from ClassiPhage and ClassiPhage 2.0 (**Figure 5**).

**Figure 5:**
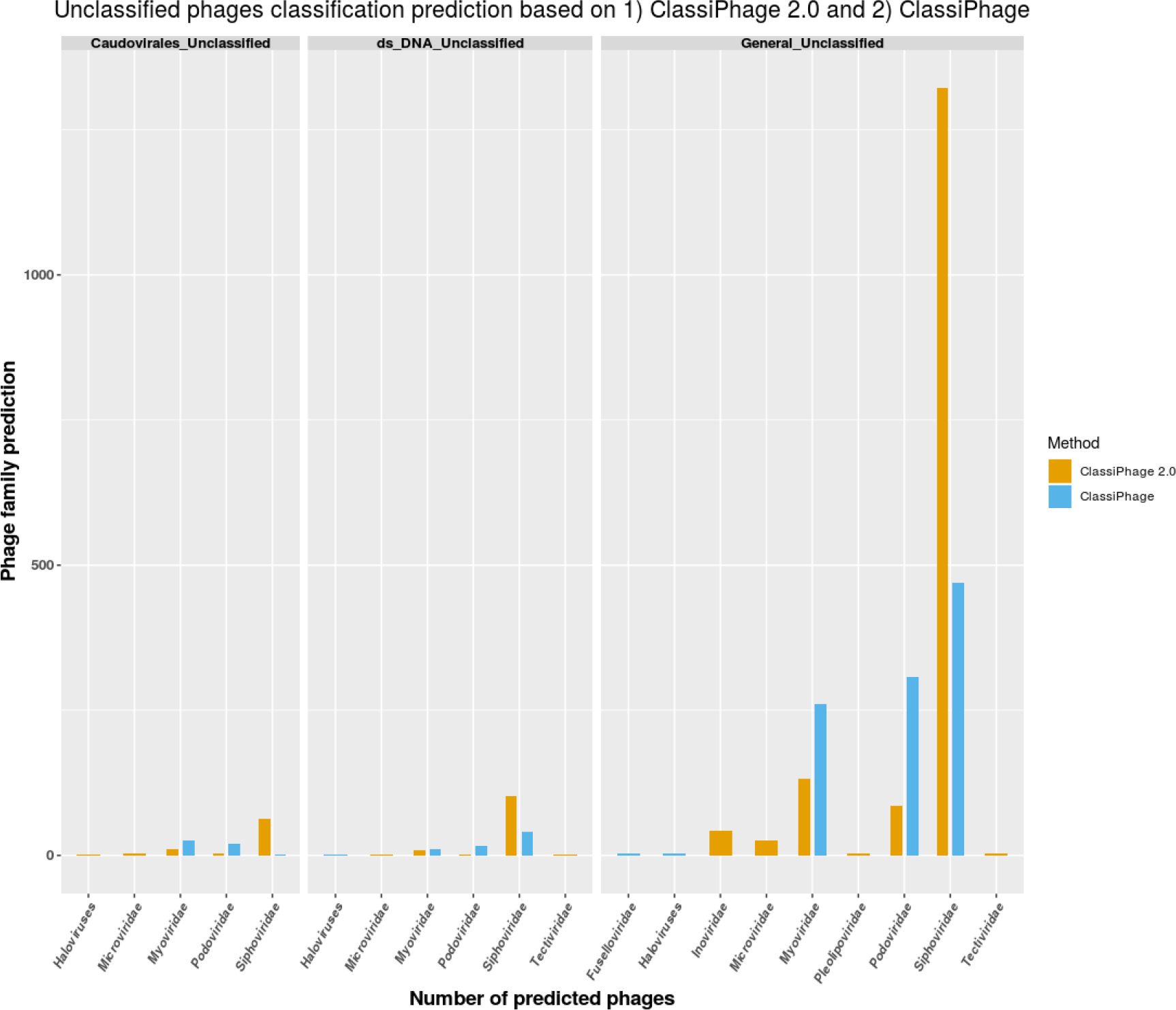
Barplot representing the classification of the unclassified phage dataset based on ClassiPhage 2.0 and ClassiPhage. A bar plot summarizing phage classification prediction of 1) ds/DNA/*Caudovirales*, 2) ds/DNA unclassified phages and 3) generally unclassified phages based on ClassiPhage 2.0 (yellow bars) and ClassiPhage (blue bars). Displaying the count number (y-axis), and the grouped phage family prediction (x-axis).

HMM based phage classification, resulted in the classification of 835 out of 1,175 generally unclassified phages into 5 of the 12 phage families (3 *Fuselloviridae*, 3 *Haloviruses*, 261 *Myoviridae*, 307 *Podoviridae* and 261 *Siphoviridae*), and resulted in the classification of 67 out of 105 ds/DNA (1 *Halovirus*,10 *Myoviridae*,16 *Podoviridae* and 40 *Siphoviridae*) and 48 out of 67 ds/DNA/*Caudovirales* (26 *Myoviridae*, 20 *Podoviridae* and 2 *Siphoviridae*) (Tables S5 and S9). The performance of ClassiPhage 2.0 prediction in comparison to HMM based phage classification was skewed towards *Siphoviridae* prediction, which is a consequence of the skewed input matrix of the ANN.

### Discussion

Phage classification based on phage sequencing data has long been a challenge, since phages have no conserved gene to place them on the tree of life (Rohwer & Edwards 2002). Although many pipelines exist for classification of prophages, these methods are based on the assumption that phages are monophyletic in origin and thus based on pairwise-alignment hits (Meier-kolthoff & Go 2018). This makes the classification of newly sequenced phages biased towards phage sequences available in the databases (Bolduc et al. 2017) and which is mostly skewed towards *Caudovirales (Skewes-cox et al. 2014)*. Therefore it is necessary to develop comprehensive computational methods for phage classification.

As stated by (Reyes & Gruber 2016), profile HMMs have an advantage over pairwise alignment in detecting remote homologs that are not part of the original MSA file used for the model’s generation. Thus profiles HMMs are more sensitive when dealing with the highly complex and diverse phages and have the potential to increase the spectrum of detectable entities. On the other hand, since HMMs rely, to some degree, on the similarity to already known sequences available in the database, and since they represent a few sequences for a few over represented viral families, means that characterizing a greater number of viral sequences and regularly updating sequence databases are crucial for this method to be effective in the future (Skewes-cox et al. 2014; Reyes et al. 2017; Reyes & Gruber 2016). Although no HMMs exist for all phage proteins, the high scoring hits to a number of HMMs derived from a phage family were enough to classify a phage based on sequence information (Chibani et al. 2019, accepted). This means that combining multiple HMM hits is crucial since no single profile HMM can assess the true viral diversity of any sequenced dataset.

To this end, we developed and applied a novel ML approach called ClassiPhage 2.0, which allows the classification of phages based on their hits into one of 12 phage families. We demonstrate that by using multiple profiles HMM as input features, derived from phage proteins out of 12 phage families, we were able to predict the phage’s taxonomic classification. Overall, we found that the method proved to be quite robust, within a range of reasonable parameter values, for the classification of the testing phage dataset, and for the assignment of a taxonomic classification of the unclassified phage dataset. However, supervised learning algorithms highly depend on the amount and quality of input data (Schmidhuber 2015). As it has been shown, phage information available in public databases is heavily biased with sequenced *Caudovirales (Skewes-cox et al. 2014; Reyes et al. 2017; Grazziotin et al. 2017)* and a large proportion of phage families are underrepresented. This further emphasizes the importance of better and more comprehensive viral databases, enriching sequence representation of each of the viral taxa, which in turn will lead to robust models constructions and thus more sensitive and comprehensive input for ML classifiers (Manavalan, Tae H. Shin, et al. 2018; Arango-Argoty et al. 2018; Amgarten et al. 2018). A misclassification resulting from this approach is due to the random split nature of k-fold cross-validation. This creates the risk for the model to predict an entry of a family that was entirely absent from its training data, due to the presence of phage families with low number of HMMs associated. As our method’s accuracy is highly dependent on the quality and accuracy of the input data, the better and more diverse the HMM models are, the better the neural network performs. That is to say that 1) whenever HMM hits are generally shared between multiple phage families such as “polymerases” or 2) if no HMM score was generated when scanning a phage proteome with the profile HMM models, then predictions are ambiguous in the first or cannot be made in the latter case. When scan outputs are not generated, the cause is that the phage belongs to a new phage family or is distant from the known phages (Roux et al. 2015). Finally, we expect the population of phage families with low abundant phages, from viral metagenomic datasets analysis. Since ANNs are known to perform better with an increasing size of a benchmark dataset (Morota et al. 2018), we foresee the improvement of ClassiPhage 2.0.

## Conclusion

In this study, we introduced a novel method which we call ClassiPhage 2.0. The method predicts a taxonomic phage family classification, resulting from multi-HMM hits of phages proteomes. We constructed ClassiPhage 2.0 using 5,920 refined profile HMMs as input features, derived from 7,342 phages classified into 12 phage families.

The results indicated that ClassiPhage 2.0 can be applied to predict a phage taxonomic classification at the family level with high accuracy. While these results are promising when observing the classification performance of one family on its own, it has proven challenging to accurately represent them in the context of all investigated families. To further elevate the performance of the neural network, as more phage data becomes available, more specific profile HMMs could be generated, improving the input datasets. In addition, the model could also be extended to include more features than HMM profile hits. This method can be further applied, for the prediction of well-delimited taxonomic groups such as subfamilies or families when profiles HMMs per subfamilies become well defined. Furthermore, the spectrum of potential applications of this approach is a general one and doesn’t have to be limited to viral classification, rather could be applied to many other classification problems in bioinformatics.

This is a tool under active development to be made available as a publicly accessible easy-to-use web service, and we envisage its growing application on a variety of forthcoming projects.

## Supplementary Data

**Supplemental Figure 1:**

**Figure S1:**
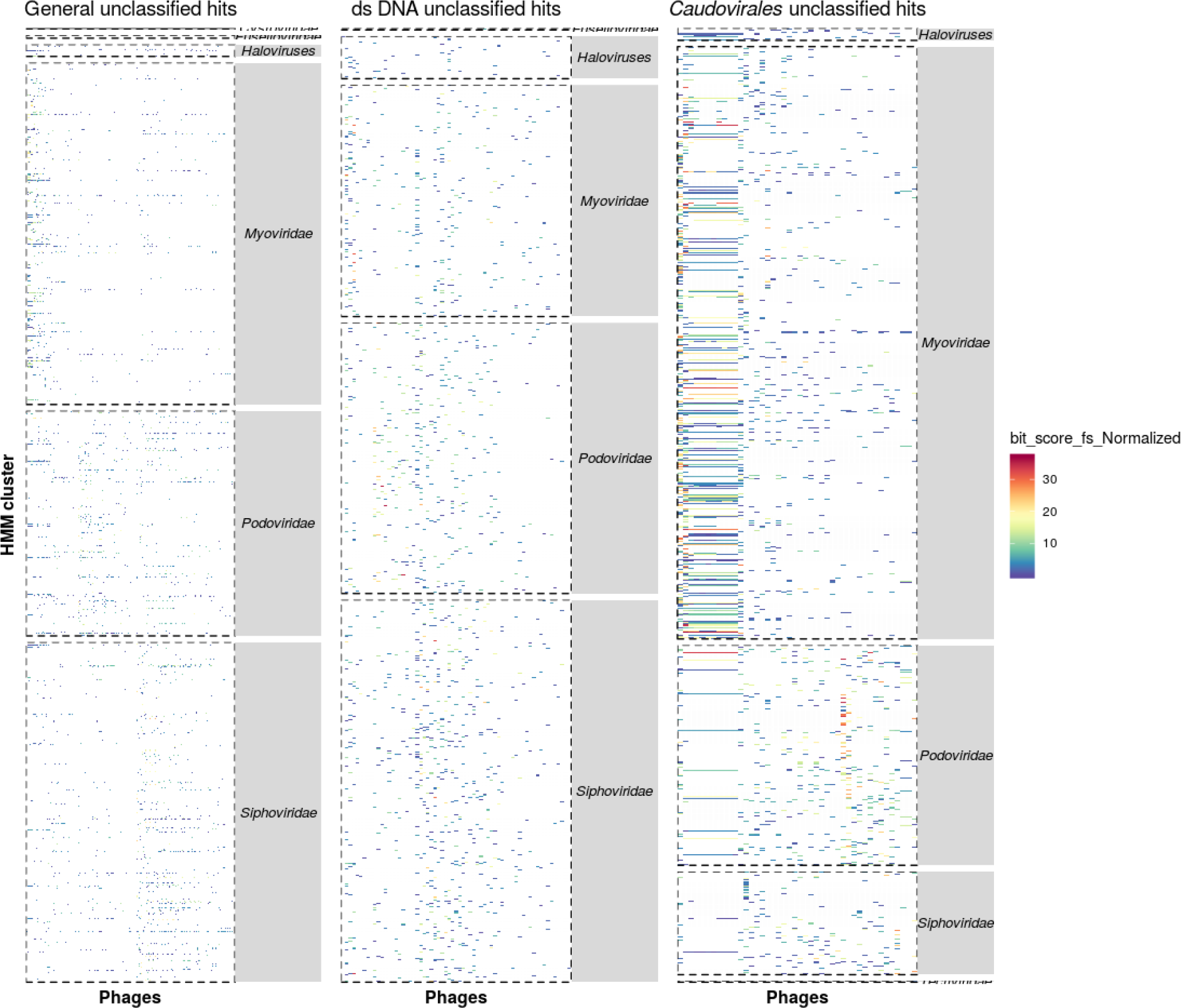
Heatmap of phage family prediction of *Caudovirales* unclassified phages depending on combination of HMM hits. The scan of the protein sequences derived from unclassified phages, was conducted by the profile HMMs of 12 phage families. The heatmap is split into 3 subplots (Generally unclassified phages, ds/DNA unclassified phages and ds/DNA/*Caudovirales*) where the phage family prediction is presented on the y-axis. The bit-score of the HMM matches was normalized by the size (in bp) of the HMM’s consensus sequence (data see Table S5). The results are color-coded from blue (low-score) to red (high-score).

**Supplemental Table S1:** All phage dataset information

Phages test dataset downloaded from the millardlab database. The table contains information for the phage, its classification and subclassification, size and accession number.

**Supplemental Table S2:** InputFamily generated HMMs scanning TargetFamily CDS Refined HMMs derived from classified phages scanning all downloaded classified phage proteomes. This table contains information for the cluster and its length, protein hit information, which phage the protein is extracted from, the phages host, the input phages classification, the scanned CDS phage classification and hmmscan information.

**Supplemental Table S3:** ClassiPhage 2.0 input matrix

Input matrix generated used as input to train and test ClassiPhage 2.0. This table contains information of the phage, its classification and bit-score values resulting from refined HMMs scan of the phage derived CDS.

**Supplemental S4:** Prediction layout of the ANN performed on the input matrix

ClassiPhage 2.0 predicted classification of classified phages. This table contains information about the phage, it’s published classification and ClassiPhage’s 2.0 classification value ranging from [0,1]. An output close to 1 is ClassiPhage’s 2.0 best predicted taxonomic classification.

**Supplemental Table S5:** InputFamily generated HMMs scanning unclassified phage CDS Refined HMMs derived from classified phages scanning all downloaded classified phage proteomes. This table contains information for the cluster and its length, protein hit information, which phage the protein is extracted from, the phages host, the input phages classification and hmmscan information.

**Supplemental Table S6:** Unclassified phage dataset matrix input for ClassiPhage 2.0

Input matrix generated used as an external dataset for classification using ClassiPhage 2.0 model. This table contains information of the phage, unknown classification tag classification and bit-score values resulting from refined HMMs scan of the phage derived CDS.

**Supplemental Table S7:** Prediction layout of the ANN for the unclassified phages dataset ClassiPhage 2.0 predicted classification of unclassified phages. This table contains information about the phage, 0 values for published classification and ClassiPhage’s 2.0 classification values ranging from [0,1]. An output close to 1 is ClassiPhage’s best predicted taxonomic classification.

**Supplemental Table S8:** Unclassified phage dataset predicted taxonomic classification via ClassiPhage 2.0 and ClassiPhages methods.

**Supplemental Table S9:** ANN prediction of unclassified Vibriophage dataset classified in Chibani et al. 2019(accepted).

Excerpt out of Table S7, which contains information about ClassiPhage 2.0 output of the same set of unclassified vibriophages classified by Chibani et al. 2019(accepted).

## Funding

KAAD for stipend, Department of Genomics and Applied Microbiology, Open access fund of DFG.

## Availability of data and materials

HMMs download available on http://appmibio.uni-goettingen.de/index.php?sec=sw (To be made public once manuscript is accepted)

## Competing interests

The authors declare that they have no competing interests.

## Author’s contributions

CC performed research, designed algorithm, performed data analysis, wrote manuscript, FM designed algorithm, wrote program, performed data analysis, AF wrote program to refine Markov Models, SD designed algorithm, HL designed research, analyzed data, wrote manuscript.

## Acknowledgements

We thank Tarek Morsi and Marc Dornieden for excellent IT-support. We thank the Goettinge Genomics Laboratory G2L for hosting. We acknowledge the support by the German research Foundation and the Open Access Fund of the Goettingen University.

## Consent for publication

Not applicable.

